# Distinct EBV-Associated Phenotypes Due to a Novel Homozygous Missense Variant in *CD27*

**DOI:** 10.64898/2026.01.15.698722

**Authors:** Rui Yang, Solomon K. Turunbedu, Sara Nandiwada, Choladda V. Curry, Tarek M. Elghetany, Brooks P. Scull, Junnan Geng, Alexander Vargas-Hernandez, Ivan K. Chinn, Sebastian Ochoa, Carl E. Allen, Lisa Forbes Satter

## Abstract

Biallelic deficiencies of CD27 and its ligand CD70 underlie selective susceptibility to Epstein-Barr virus (EBV) infection and its acute and chronic complications, underscoring their non-redundant roles in anti-EBV immunity. To date, 16 pathogenic *CD27* variants have been reported. Here, we describe three patients from two unrelated families, homozygous for a novel loss-of-function (LOF) *CD27* variant, resulting in substitution of serine 70 with proline (S70P). All three patients presented with EBV viremia and lymphoproliferative disease, with variable immune dysregulation or recurrent otosinopulmonary infections. One patient developed EBV-associated Hodgkin lymphoma. Functional studies demonstrated that the S70P variant impaired surface expression of CD27 and abolished CD70 binding, rendering complete LOF. Together, S70P represents a novel pathogenic *CD27* variant causing autosomal recessive (AR) CD27 deficiency, characterized by a unified susceptibility to EBV yet variable clinical manifestations, ranging from chronic viremia to malignancy.

## INTRODUCTION

Epstein-Barr virus (EBV) infects over 90% of adults worldwide (Zamora, 2011). Seroprevalence rises with age, reaching ∼30% by ages 1-5 years, 40-55% at 6-9 years, 50-65% at 10-14 years, and 70-80% by 15-19 years (Balfour et al., 2013b; Condon et al., 2014; Dowd et al., 2013). Most primary infections are asymptomatic, while symptomatic cases usually present as mild infectious mononucleosis (IM) (Balfour et al., 2013b; Crawford et al., 2006; Topp et al., 2015). While most EBV infections follow a subclinical or mild course, severe disease can occur, including fulminant presentations (meningoencephalitis, airway obstruction, fulminant hepatitis), immune dysregulation (hemophagocytic lymphohistiocytosis, HLH; refractory cytopenias), chronic infections (persistent EBV viremia, chronic active EBV–CAEBV), lymphoproliferative disease (LPD), and EBV-related malignancies (Balfour et al., 2013a; Bollard and Cohen, 2018; Kawada et al., 2023; Lino and Ghosh, 2021; Sacco et al., 2023; Tangye et al., 2017). The marked inter-individual variability in susceptibility remains poorly understood. Twin studies reveal a strong inheritable component to both IM and Hodgkin lymphoma (Mack et al., 1995; Rostgaard et al., 2014), and discoveries of inborn errors of immunity (IEIs) with selective susceptibility to EBV demonstrate that EBV susceptibility, at least in some patients, follows a monogenic basis (Deau et al., 2014; Elkaim et al., 2016; Latour, 2025; Lucas et al., 2014; Martin et al., 2024; Tangye, 2020).

IEIs predisposing to EBV can be categorized as either part of broader infection susceptibility or as selective EBV susceptibility. The former includes autosomal dominant (AD) *GATA2* deficiency, X-linked recessive (XLR) *WAS* deficiency, and AR deficiencies of *STIM1*, *SLP76*, *STK4*, *DOCK8*, and *CORO1A*, as well as activated phosphoinositide 3-kinase δ syndrome 1 and 2 (APDS1 and APDS2), caused by pathogenic *PIK3CD* and *PIK3R1* variants. These disorders compromise T-cell activation or effector function, often in combination with other immunological defects (Latour, 2025). In contrast, twelve forms of IEIs predisposing to selective EBV susceptibility, defined by the prevalence of severe EBV disease or the predominance of EBV infection as the key clinical manifestation, often accompanied by variable degrees of other infections or immune dysregulation, have been reported (Deau et al., 2014; Elkaim et al., 2016; Latour, 2025; Lucas et al., 2014; Martin et al., 2024; Tangye, 2020). X-linked lymphoproliferative disorder 1 (XLP1) and X-linked immunodeficiency with magnesium defect, EBV infection, and neoplasia (X-MEN) syndrome, caused by XLR *SH2D1A* and *MAGT1* deficiencies, respectively, both disrupt T- and NK-cell activation and cytotoxic effector function (Latour, 2025). AR deficiencies of *CTPS1*, *RASGRP1*, *ITK*, and *CARMIL2* impair the T-cell receptor (TCR) signaling cascade. AR *IL27RA* and XLR *XIAP* deficiencies limit the expansion and survival of EBV-specific effector T cells (Latour, 2025; Martin et al., 2024). AR deficiencies of CD137L–CD137 and CD70–CD27 axes, two TNF receptor superfamily ligand–receptor pairs, impair the priming, activation, and effector function of EBV-specific T cells (Latour, 2025). Among all twelve IEIs with selective EBV susceptibility, AR CD27 and CD70 deficiencies confer the highest malignancy risk, with EBV-driven lymphomas reported in 16 of 38 and 10 of 17 patients, respectively, resulting in substantial mortality and morbidity (Baskin et al., 2024; Ghosh et al., 2020; Golchehre et al., 2023; Köse et al., 2022). Here, we identify three patients with AR CD27 deficiency and characterize their clinical, genetic, molecular, histologic, and immunologic features, thereby expanding the spectrum of EBV susceptibility.

## RESULTS

### Identification of three patients with EBV-associated disease

We studied three patients from two unrelated families with EBV-associated disease. Patient P1 (**Fig. 1A, Kindred A**), a girl born to nonconsanguineous Afro-Latino parents of Honduran origin, presented at 3 years of age with a right neck mass (**Fig. 1 – S1A** and **S1B, Table 1**). An excisional lymph node (LN) biopsy demonstrated polymorphous infiltrates with frequent large atypical Hodgkin Reed-Sternberg (HRS) cells surrounded by thick fibrous septae (**Fig. 1B**). By immunohistochemistry (IHC), HRS cells were CD3^−^, partial CD20^+^, CD45^−^, CD30^+^, and PAX5^dim^, consistent with nodular sclerosis Hodgkin lymphoma (HL) (**Fig. 1 – S2A – S2C** and **Fig. 1C**). Notably, both EBV-encoded small RNAs (EBER) by *in situ* hybridization (ISH) and latent membrane protein 1 (LMP1) by IHC were positive, suggesting an EBV-driven etiology (**Fig. 1D**). Despite four cycles of doxorubicin, bleomycin, vincristine, etoposide, prednisone, and cyclophosphamide (ABVE-PC) chemotherapy, EBV viremia persisted, and a repeat LN biopsy showed EBER-positive immunoblasts (**Fig. 1 – S1B** and **Fig. 1 – S2D and S2E**). She subsequently received B-cell depletion therapy with rituximab, which initially cleared the viremia, but recurrence followed. This prompted the decision to pursue hematopoietic stem cell transplantation (HSCT). At 4 years of age, she underwent a haploidentical HSCT from her father, following conditioning with fludarabine, alemtuzumab, and total body irradiation (TBI). Her post-transplant course was complicated by CMV viremia, for which she received adoptive T cell therapy (Hanley et al., 2009), hepatic graft-versus-host disease (GVHD), and *Enterobacter* and *Serratia* bacteremia (**Fig. 1 – S1B**). Due to declining mixed chimerism and borderline EBV viremia, she underwent a second haploidentical HSCT from her father at age 5. Thereafter, EBV viremia resolved and has remained negative since age 7, despite a persistently low mixed chimerism of only approximately 20% donor cells. At age 13, she developed decompensated heart failure with severe biventricular dysfunction, likely secondary prior cardiotoxic chemotherapy, and since age 14 has been supported with a ventricular assistance device (VAD) (**Fig. 1 – S1B**).

**Figure 1.**
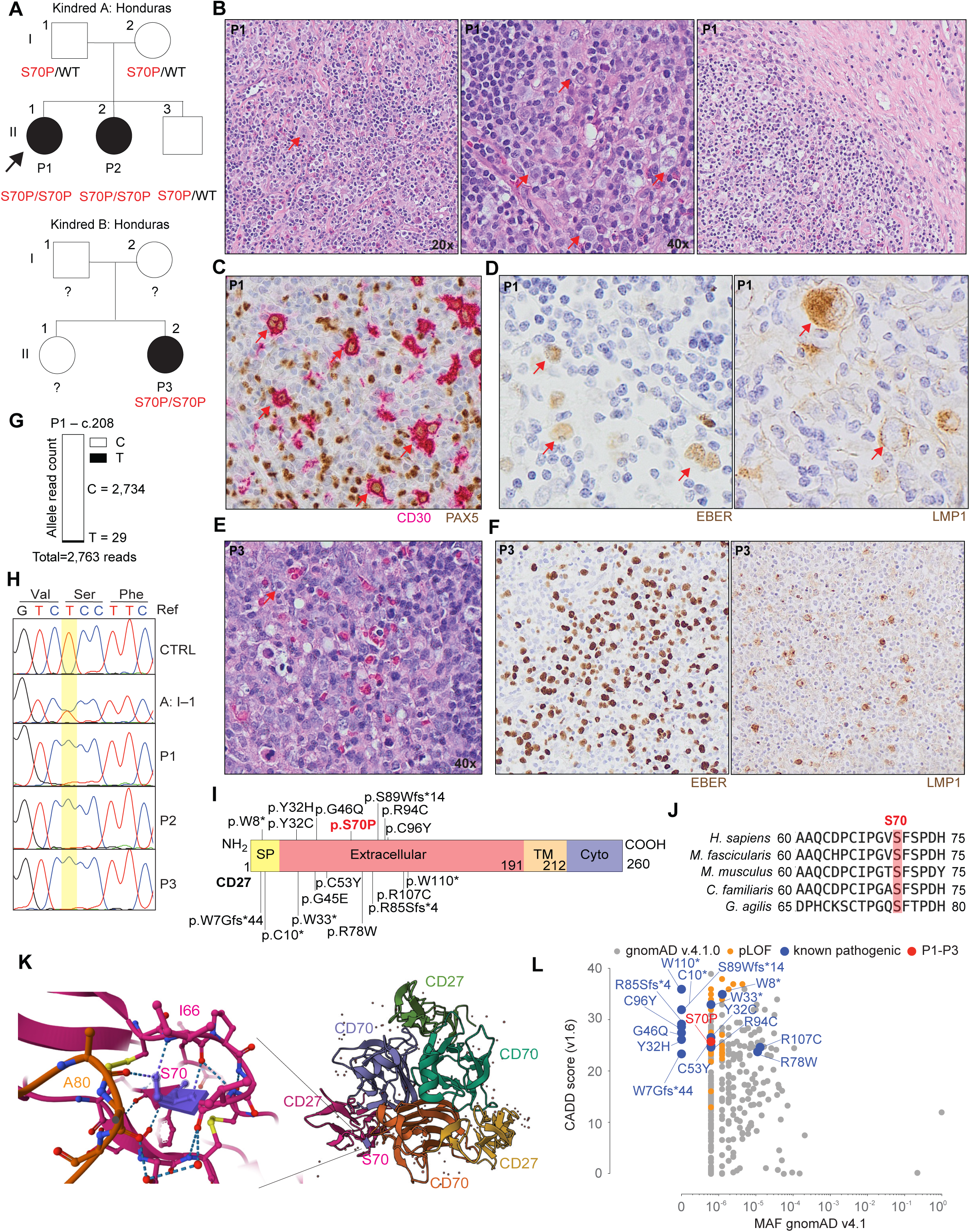
Three patients with inherited CD27 deficiency due to a homozygous S70P variant. **(A)** Pedigrees showing familial segregation of *CD27* S70P variant. **(B)** H&E images of a lymph node biopsy from P1. **(C)** Multiplex immunohistochemistry of PAX5 and CD30 on a lymph node biopsy from P1. **(D)** In situ hybridization for EBER and immunohistochemistry for LMP1 on a lymph node biopsy from P1. **(E)** High-power H&E image (40x) of a lymph node biopsy from P3. **(F)** In situ hybridization for EBER and immunohistochemistry for LMP1 on a lymph node biopsy from P3. **(G)** Sanger sequencing showing variant allele frequency of S70P *CD27* variant in post-transplant DNA from P1. **(H)** Sanger sequencing confirming the homozygous S70P *CD27* variant in P1, P2, and P3 as well as their family members. **(I)** Schematic representation of CD27 protein domains with the location of the indicated variants. **(J)** Conservation of the Ser70 residue across CD27 orthologs from multiple species. (**K**) Three-dimensional crystal structure of the CD27 trimer in complex with CD70 trimer, highlighting the position of S70P variant. **(L)** Scatter plot of CADD v1.6 scores against minor allele frequencies (MAFs) of *CD27* variants, showing S70P variant, known pathogenic variants, predicted LOF variants, along with homozygous and heterozygous variants from gnomAD v4.1.

**Table 1:** Clinical summary of three patients with inherited CD27 deficiency.

|  | **P1** | **P2** | **P3** |
| --- | --- | --- | --- |
| **Origin** | Latino | Latino | Latino |
| **Age of onset (years)** | 3 | 6 | 2 |
| **Initial presentation** | Neck mass, diagnosed with Hodgkin's Lymphoma | Pneumonia, pleural effusion, and pancytopenia | Neck swelling, complicated by airway obstruction |
| **EBV viremia on presentation** | Yes | Yes | Yes |
| **Blood EBV level on presentation (units lab specific)** | 122,043 (copies per µg DNA from blood) (0-200) | > 500,000 IU/mL (< 111 IU/mL, one IU is equal to 9.03 copies of EBV) | 249,204 (copies per µg DNA from blood) (0-200) |
| **Hemophagocytic lymphohistiocytosis (HLH)** | No | Yes | No |
| **Chronic EBV viremia** | Yes | Yes | Yes |
| **Lymphoproliferative disease (LPD)** | Yes | Yes | Yes |
| **Malignancy** | Yes, Hodgkin Lymphoma | No | No |
| **recurrent otosinopulmonary infection** | Yes, mostly during post-HSCT peroid | Yes, confounded by prematurity and bronchopulmonary dysplasia | Yes |
| **Immunoglobulin replacement therapy (IgRT) dependence** | No | No | No |
| **Hematopoietic stem cell transplantation (HSCT)** | Yes, twice (2nd HSCT due to persistent EBV viremia and low donor chimerism) | Yes | No |
| **Age at HSCT (years)** | 1st: 4<br>2nd: 5 | 7 | NA |
| **Conditioning regimen** | Fludarabine, Campath, and TBI | Fludarabine, cyclophosphamide, TBI, and post-HSCT cyclophosphamide | NA |
| **HSCT donor** | Haploidentical (father) | Haploidentical (mother) | NA |
| **Chimerism at last follow up** | ~20% donor | 100% donor | NA |
| **Age at last follow up (years)** | 15 | 12 | 11 |
| **Age at EBV viremia resolution (years)** | 6 | 7 | EBV viremia persists |

Patient P2, sister of P1 (**Fig. 1A, Kindred A, Table 1**), was born at 25 weeks’ gestation because of recurrent fetal heart rate decelerations. In early childhood from ages 1 to 6, she experienced recurrent upper and lower respiratory infections (URTIs and LRTIs) in the setting of bronchopulmonary dysplasia and pulmonary hypertension related to extreme prematurity. At age 6, she was hospitalized with respiratory insufficiency due to left lower lobe (LLL) pneumonia and pleural effusion, accompanied by pancytopenia and profound EBV viremia (**Fig. 1 – S1B** and **Table 1**). Over the following month, she developed progressive diffuse lymphadenopathy and hepatosplenomegaly. Excisional biopsy of a cervical LN revealed EBER-positive interfollicular immunoblastic expansion. She was diagnosed with CAEBV and secondary HLH. Despite treatment with rituximab and the HLH-94 regimen (Henter et al., 2002), her EBV viremia persisted (**Fig. 1 – S1B** and **Table 1**), which prompted the decision to proceed with a haploidentical HSCT from her mother at age 7, using conditioning with fludarabine, cyclophosphamide, and TBI, following bridging with the HLH-94 regimen. Her post-transplant course was complicated by *Stenotrophomonas* and *Achromobacter* bacteremia and hemorrhagic cystitis. Since HSCT, her blood EBV levels have remained undetectable. She experienced one episode each of bacteremia due to *Streptococcus pneumoniae* and methicillin-resistant *Staphylococcus aureus* (MRSA) at age 7, but has remained otherwise healthy since. Both parents and the younger brother of P1 and P2 have remained healthy.

Patient P3 (**Fig. 1A**, **Kindred B** and **Table 1**), a girl born to nonconsanguineous Afro-Latino parents of Honduran origin, presented with fever and a right neck mass at 3 years of age, followed by progressive worsening of cervical lymphadenopathy that compromised her airway, requiring endotracheal intubation for airway protection. Excisional biopsy of a cervical LN revealed intact follicular structures but marked paracortical expansion by a heterogeneous population of lymphocytes, histiocytes, plasma cells, and immunoblasts (**Fig. 1E** and **Fig. 1 – S2F – S2H**). Parafollicular infiltrates, including immunoblasts, were scatteredly positive for EBER ISH and LMP1 IHC (**Fig. 1F**). Her blood EBV levels were markedly elevated (**Fig. 1 – S1B** and **Table 1**). She underwent rituximab induction, after which she has remained asymptomatic despite persistently elevated EBV viremia (**Fig. 1 – S1A**). An older sister and both parents have remained healthy.

### Identification of a rare missense *CD27* variant in these patients

P2 and P3 underwent whole exome sequencing (WES). Both parents and the younger brother of kindred A were also sequenced, enabling trio-WES analysis. P2 and P3 shared the same homozygous single-nucleotide variant (SNV) in *CD27*, c.208T>C, resulting in a missense substitution (p.Ser70Pro;S70P) (**Fig. 1A**). P1 underwent transplantation before WES was clinically available. However, amplicon sequencing of P1’s saliva-derived DNA revealed that 99% (2,734/2,764) of the reads harbored mutant c.208C, with the remaining 1% of reads (29/2,764) being WT c.208T (**Fig. 1G**), and sanger sequencing using post-transplant saliva-derived DNA also showed homozygosity for the mutant allele (**Fig. 1H**). Collectively, these data suggested that P1 was homozygous for c.208T>C, with the low-level WT signal most consistent with minor donor-derived DNA contribution in saliva (Garbieri et al., 2017), given poor donor chimerism and haploidentical transplantation. Sanger sequencing in P2 and P3 also confirmed the T-to-C substitution at c.208, consistent with the S70P missense substitution (**Fig. 1H**). Both parents and the younger brother of P1 and P2 were heterozygous for the variant, whereas the genotypes of the P3 family members were not obtained (**Fig. 1H**). The Ser70 residue localized to the extracellular domain of CD27 protein, where nearly all disease-causing CD27 variants reported to date are clustered, except for three (p.W7Gfs*44, p.W8*, and p.C10*) that reside in the signal peptide of the precursor protein (Ghosh et al., 2020) (**Fig. 1I**). Ser70 is evolutionarily conserved across multiple mammalian species (**Fig. 1J**). Ser70 and Phe71 form a short β-strand flanked by coil regions (**Fig. 1K**). Notably, Ser70 contributes to CD27 stability through two hydrogen bonds with Ile66, and further interacts across the CD27-CD70 interface by forming a hydrogen bond with Ala80 and a van der Waals contact with Leu81 of CD70 (**Fig. 1K**) (Liu et al., 2021). Thus, we identified a homozygous S70P variant in all three affected patients, which may underlie their disease phenotype.

### S70P variant is likely deleterious based on bioinformatic predictions

Like all other known pathogenic *CD27* variants, S70P was rare in the general population. It was absent from gnomAD v2.1 but present in BRAVO with a minor allele frequency (MAF) of 0.0000298 and in gnomAD v4.1 with an overall MAF of 0.00000682 (Chen et al., 2022; Karczewski et al., 2020; Taliun et al., 2021). Notably, the highest MAF was in the Admixed American population of gnomAD v4.1, where it reached 0.00015 (**Fig. 1L**). No homozygotes were identified in BRAVO, gnomAD v2.1, 3.1, or 4.1 (Chen et al., 2022; Karczewski et al., 2020; Taliun et al., 2021). The combined MAF for all known pathogenic variants, including the newly identified S70P, together with predicted LOF (pLOF) variants, was approximately 0.000056, corresponding to an estimated biallelic prevalence of < 1 in 100 million. The Combined Annotation Dependent Depletion (CADD) v1.6 PHRED score for the S70P variant was high at 25.8, placing it among the top ∼0.26% most deleterious variants genome-wide (Kircher et al., 2014; Rentzsch et al., 2021; Zhang et al., 2018). Similarly elevated CADD scores were observed for all disease-causing *CD27* variants (**Fig. 1L**). PolyPhen-2 and AlphaMissense scores were high at 0.999 and 0.9612 for S70P, respectively, both consistent with deleteriousness (Adzhubei et al., 2013; Adzhubei et al., 2010; Cheng et al., 2023). Thus, bioinformatic prediction tools suggest deleteriousness and evidence from population genetics does not contradict our genetic hypothesis.

### S70P is LOF

We investigated the functional impact of the S70P variant using a balanced overexpression system, in which only one copy of the transgene was integrated and expressed under the same tetracycline-inducible promoter (Kamath and Matreyek, 2024; Matreyek et al., 2018; Shukla et al., 2025). Overexpression of wild-type (WT) *CD27* resulted in robust CD27 surface expression, detected by two distinct monoclonal antibodies (mAbs) (**Fig. 2A**). In contrast, overexpression of an unrelated control protein ACE2 or a known loss-of-expression (LOE) variant, W33*, abolished CD27 surface expression (**Fig. 2A**) (Ghosh et al., 2020). Overexpression of the S70P variant exhibited only minimal surface expression, detectable by both mAbs, significantly lower than WT but slightly higher than W33* and ACE2 controls (**Fig. 2A**). This result suggests that the S70P variant destabilized the CD27 protein, consistent with crystallographic evidence that Ser70 forms key hydrogen bonds with Ile66 (Liu et al., 2021). We next assessed CD70 binding. Overexpression of WT, A59T, and P138L *CD27*, the two most common single nucleotide polymorphisms (SNPs) in the general population, exhibited strong, concentration-dependent binding of soluble CD70 (sCD70) binding (**Fig. 2B** and **2C**). As expected, W33* completely failed to bind sCD70 (**Fig. 2B** and **2C**). Previously reported pathogenic variants, R107C and Y32H, behaved as severely impaired hypomorphs, with markedly reduced but still detectable sCD70 binding (**Fig. 2B** and **2C**) (Alkhairy et al., 2015; Ghosh et al., 2020). In contrast, overexpression of S70P abolished sCD70 binding (**Fig. 2B** and **2C**). Pre-transplantation whole blood from P2 and P3 was examined for surface CD27 expression. CD27 expression on total lymphocytes was abolished in P2 and P3 compared with those of healthy donors (**Fig. 2D**). P1’s whole blood also showed abolished CD27 expression by clinical immunophenotyping. Thus, S70P impairs CD27 surface expression and abolishes CD70 interaction, establishing it as LOF when overexpressed, whereas endogenous CD27 expression is absent in patients homozygous for S70P.

**Figure 2.**
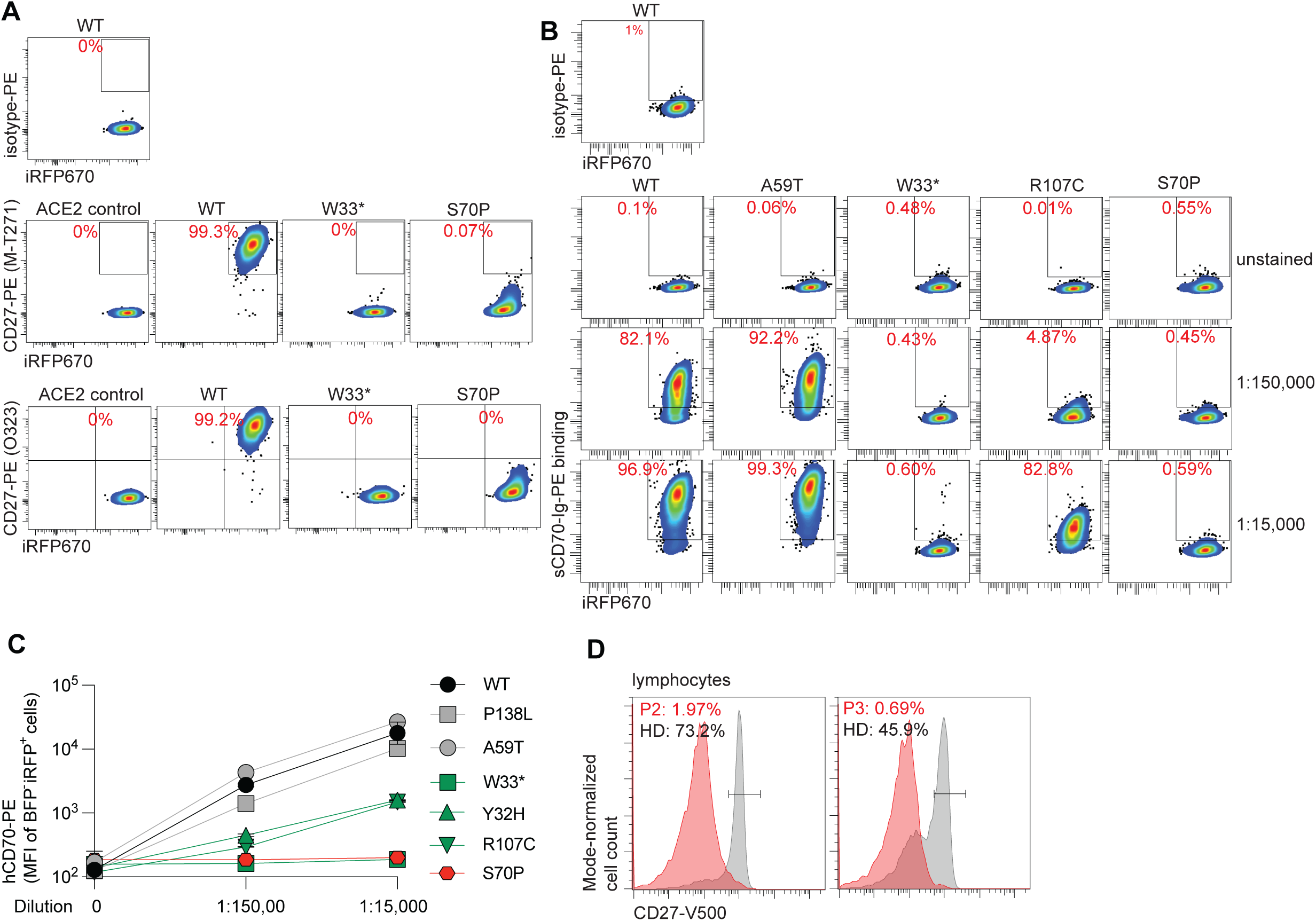
The S70P *CD27* variant is LOF. **(A)** Expression of WT or mutant *CD27* alleles on landing pad cells recombined using the Bxb1-AttB system. **(B)** Soluble CD70 binding on landing pad cells overexpressing WT or mutant *CD27* alleles. **(C)** Soluble CD70-PE mean fluorescence intensity (MFI) values plotted as dose-reponse curves for WT and mutant *CD27* alleles. **(D)** Endogenous surface CD27 expression on lymphocyte surface in healthy donors (HDs) and from patients P2 and P3.

### B cell development and antibody production are altered in S70P homozygosity

CD27 is a memory marker in B cells, and CD27 ligation promotes immunoglobulin production, B-cell priming, proliferation, and differentiation during germinal center responses (Grimsholm, 2023; Tangye et al., 1998). We investigated B cell development in patients with inherited CD27 deficiency due to the homozygous S70P variant using pre-transplantation blood samples. B cell counts were normal in P1 but below the 10^th^ percentile for age for P2 and P3 at presentation, with recovery over time in P3 (**Fig. 3A**) (Shearer et al., 2014). All three patients presented with pan-hypergammaglobulinemia, with markedly elevated IgG, IgA, and IgM (**Fig. 3B – D**), which can be seen in primary EBV infection (Andropoulos, 2012; Lo et al., 2013). P1’s IgG level remained elevated until HSCT, whereas P3’s IgG normalized by age 4 following rituximab (**Fig. 3B**). IgA and IgM levels in P1 and P3 normalized after chemotherapy and rituximab, respectively (**Fig. 3C** and **D**). Anti-tetanus and anti-diphtheria IgG levels in all three patients were protective (>= 0.1 IU/mL) (**Fig. 3E**) (Orange et al., 2012). In contrast, humoral immunity to polysaccharide antigens appeared impaired. In P3, despite completion of the 13-valent pneumococcal protein-conjugated vaccine (PCV13) primary series, only 17.4% of 13 serotypes reached 0.35 µg/mL, the threshold for protection against invasive pneumococcal disease (Orange et al., 2012; Zuzolo et al., 2025) (**Fig. 3F**). Following the 23-valent pneumococcal polysaccharide vaccine (PPSV23) at age 8, only 12/23 serotypes met criteria for response, defined as >= 1.3 µg/mL or >= 2-fold increase if pre-PPSV23 titers already exceeded 1.3 µg/mL (Orange et al., 2012; Zuzolo et al., 2025) (**Fig. 3F**). Only 6/11 non-PCV13 serotypes responded, a rate below the >= 70% expected for age (Orange et al., 2012; Zuzolo et al., 2025) (**Fig. 3G**). Although none of the patients required long-term immunoglobulin replacement therapy (IgRT), each experienced at least two episodes of severe bacterial infection, such as sepsis or pneumonia. Lastly, CD27 surface expression was markedly reduced on the IgD-negative (IgD^−^) B-cell subset in P3, compared with a healthy donor (**Fig. 3H**). Collectively, homozygous S70P ablates CD27 expression in B cells. While B cell counts are variable, antibody response to protein antigens are preserved, whereas responses to polysaccharide antigens appear impaired.

**Figure 3.**
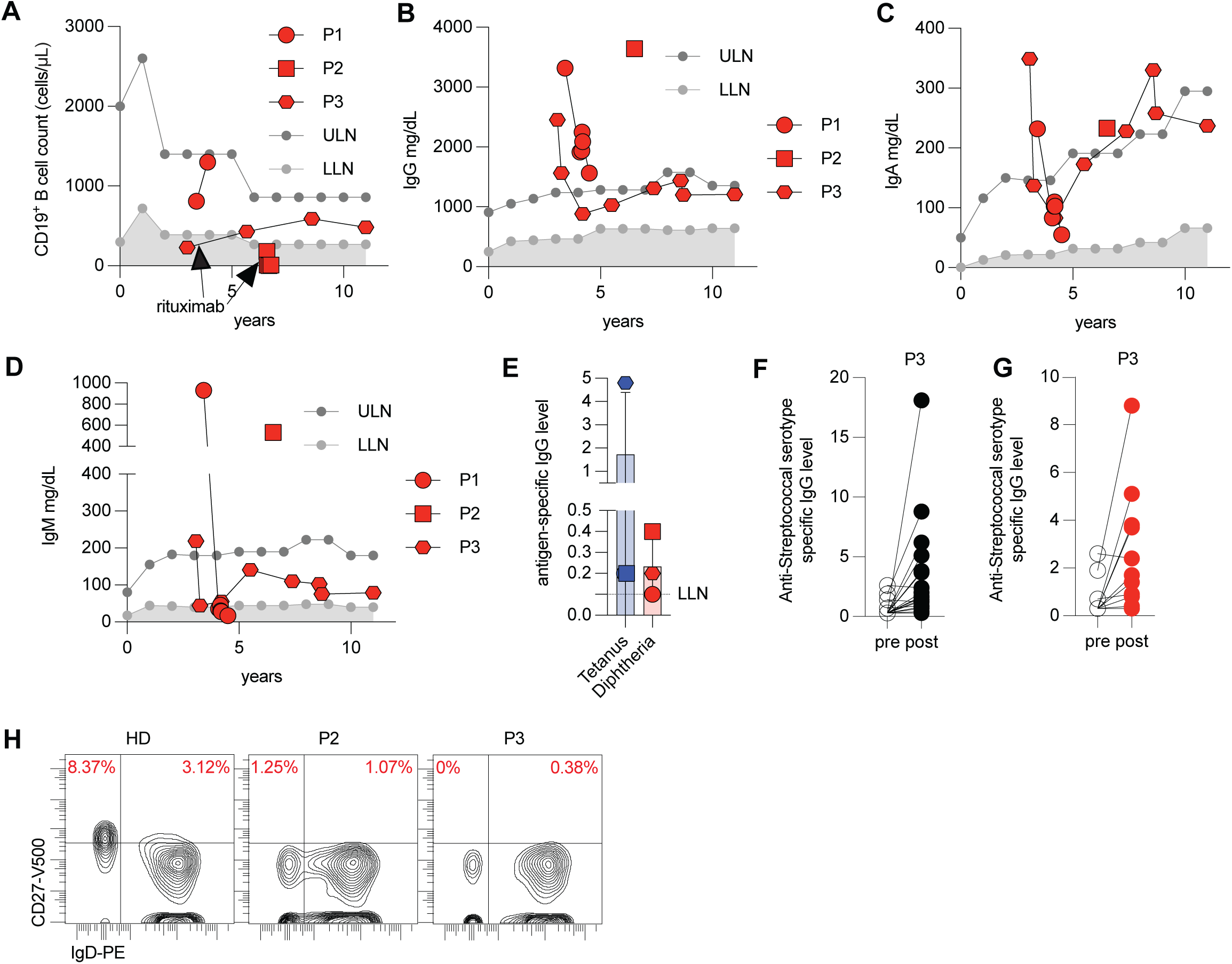
Humoral immunity is altered in inherited CD27 deficiency. **(A)** Longitudinal peripheral B cell counts in patients P1, P2, and P3. **(B – D)** Longitudinal serum immunoglobulin levels for IgG (B), IgA (C), and IgM (D). **(E)** Anti-tetanus and anti-diphtheria IgG levels in P1, P2, and P3. **(F** and **G)** Pre-and post-PPSV23 pneumococcal antibody responses: across all 23 serotypes (F) and polysaccharide-specific responses to serotypes unique to PPSV23 but not included in the Prevnar 13 vaccine (G). **(H)** Representative contour plots showing CD27 and IgD expression on CD19^+^CD20^+^ B cells.

### Absent CD27 expression on NK cells

We next studied natural killer (NK) cells, another key effector population in anti-EBV immunity in humans (Latour, 2025), using pre-transplantation blood samples. NK cell counts were normal in P1 and P3 but reduced in P2 (**Fig. 4A**). The proportions of CD56^bright^ and CD16^+^ cells were comparable to healthy donors (**Fig. 4B** and **4C**). To assess cytotoxic effector function, we measured spontaneous cytotoxicity against target K562 cells. NK cells from healthy donors, P2, or P3 were incubated with K562 cells at graded effector:target ratios in the absence or presence of IL-2. Both P2 and P3 NK cells displayed cytotoxic activity within or above the lower limit of the normal range, both with or without IL-2 (**Fig. 4D**). In humans, CD27^+^ NK cells represent a transitional subset specialized for immune regulation through cytokine production rather than direct cytotoxicity (Silva et al., 2008). As expected, CD27^+^ NK cells were proportionally more enriched in CD56^bright^ NK cells in healthy donors (**Fig. 4E**). However, CD27^+^ NK cells were depleted in P2 and P3 (**Fig. 4E**). Thus, homozygous S70P depletes CD27 expression on NK cells but preserves major subsets of NK cells and their spontaneous cytotoxic activity.

**Figure 4.**
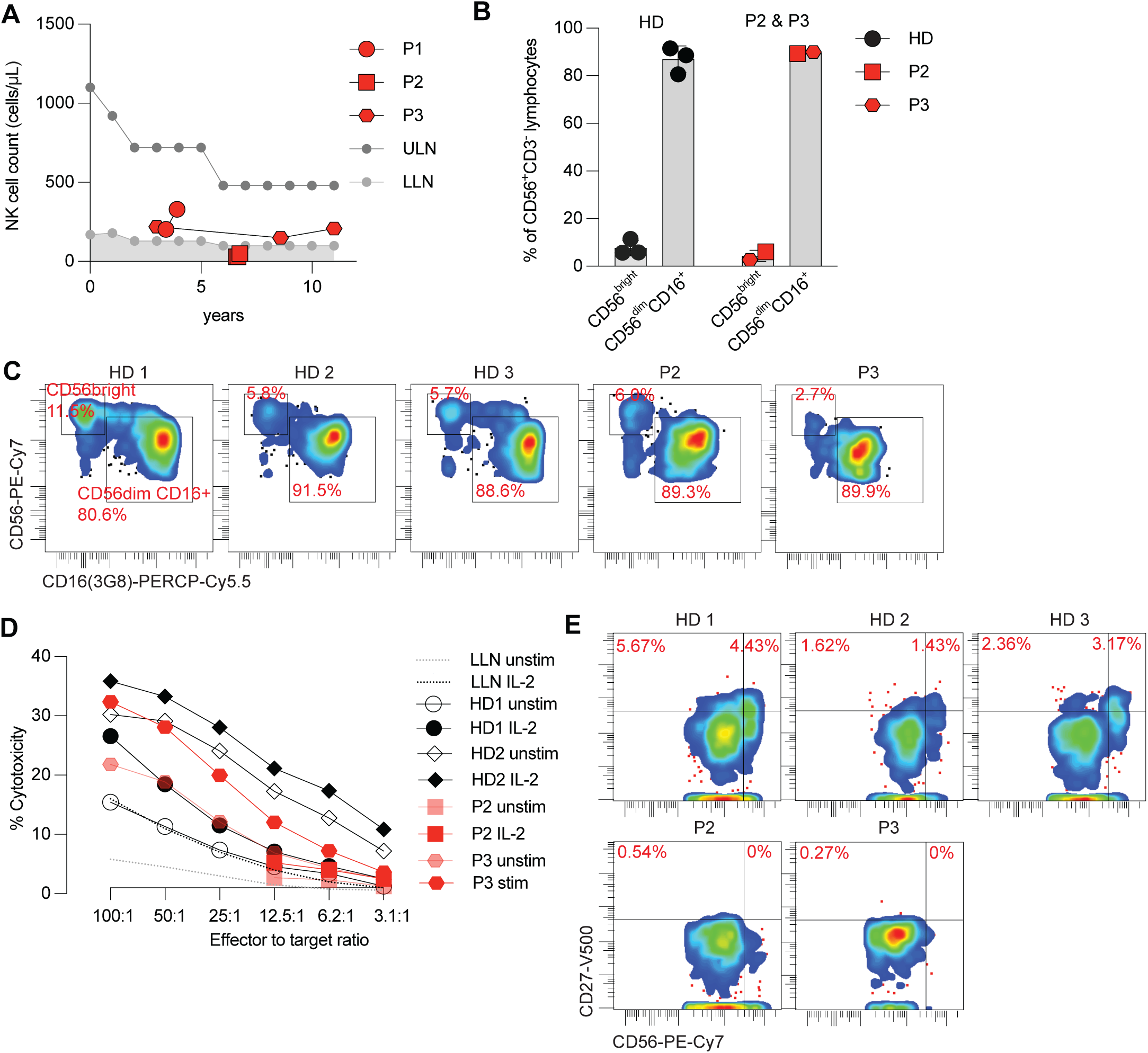
Abolished CD27 expression on NK cells. **(A)** Longitudinal peripheral NK cell counts. **(B)** Proportions of CD56^bright^ and CD56^dim^CD16^+^ NK subsets within total NK cells. **(C)** Expression of CD56 and CD16 on NK cells from HDs or patients P2 and P3. **(D)** NK cell spontaneous cytotoxic activity against K562 target cells in the presence or absence of IL-2 stimulation. **(E)** Expression of CD27 and CD56 on NK cells from HDs and from patients P2 and P3.

### Variable T-cell lymphopenia but sufficient T cell proliferation in inherited CD27 deficiency

CD70 engagement with CD27 provides a key co-stimulatory signal priming T cells for anti-EBV immunity (Ghosh et al., 2020; Latour, 2025). We therefore investigated the pre-HSCT T-cell compartment in all three patients. CD3^+^ T cell counts were normal in P3 but reduced in P1 and P2 (**Fig. 5A**). Similar patterns of CD4^+^ and CD8^+^ T-cell lymphopenia were observed in P1 and P2 (**Fig. 5B** and **5C**). CD4^+^ T cells were skewed toward a memory phenotype. The frequency of CD45RA^+^ cells was reduced in P3 and only marginally normal in P1, whereas CD45RO^+^ cells were enriched in both P1 and P3 (**Fig. 5D** and **5E**). We next assessed T-cell proliferation. T cells from all three patients exhibited normal responses to phytohemagglutinin (PHA), concanavalin A (con A), and pokeweed mitogen (PWM) (**Fig. 5F – 5H**). In contrast, antigen-driven proliferation revealed subtle defects. P2 showed a suboptimal response to tetanus, and P3 showed suboptimal responses to both tetanus and diphtheria. In contrast, T cells from all three patients proliferated robustly to *Candida* (**Fig. 5I – 5K**). Collectively, despite variable T-cell lymphopenia and occasional suboptimal responses to antigenic stimulation, overall T-cell proliferative capacity is preserved in inherited CD27 deficiency due to the homozygous S70P variant, consistent with the lack of opportunistic infections outside the peri-HSCT period.

**Figure 5.**
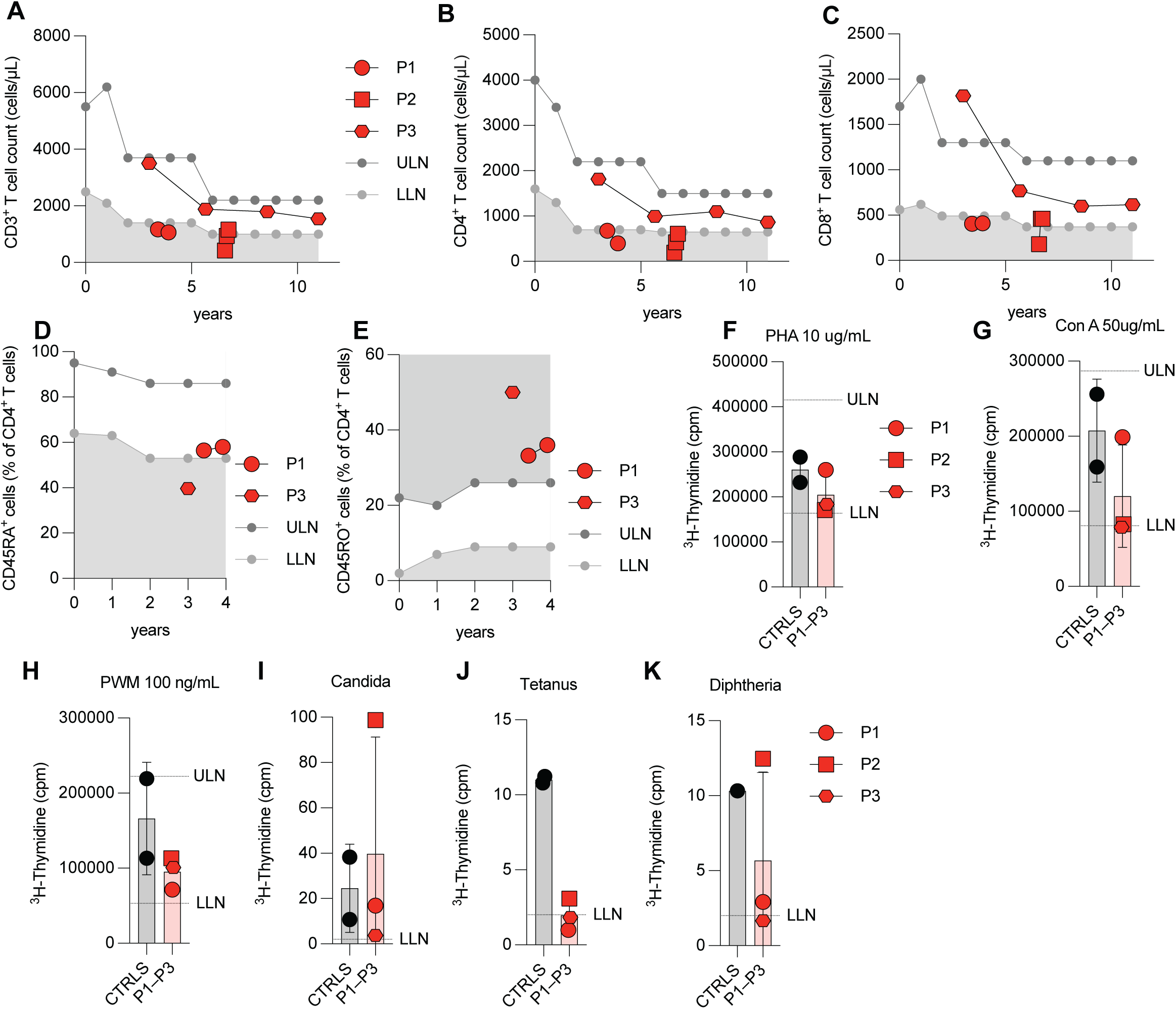
Borderline T cell lymphopenia and memory T cell skewing in inherited CD27 deficiency. **(A – C)** Longitudinal peripheral counts of total CD3^+^ T (A), CD4^+^ T (B), and CD8^+^ T lymphocytes (C). **(D and E)** Proportions of CD45RA^+^ (D) and CD45RO^+^ (E) subsets within CD4^+^ T cells. (**F – K**) T cell proliferation, measured by ^3^H-thymidine incorporation, following stimulation with PHA (F), Con A (G), PWM (H), *C.albicans* antigen (H), tetanus toxoid (J), and diphtheria toxoid (K).

### Endogenous S70P CD27 is LOF in T cells

We next investigated CD27 expression on T cells from P2 and P3. Compared with healthy donors, CD27 surface expression on T lymphocytes from P2 and P3 was markedly reduced *ex vivo* (**Fig. 6A**). T-cell blasts derived from pre-HSCT peripheral blood mononuclear cells (PBMCs) were analyzed for CD27 expression following CD2, CD3, and CD28 stimulation. In a healthy donor, both CD4^+^ and CD8^+^ T cells rapidly upregulated CD27 surface expression, which subsequently declined as cells returned to quiescence. A major subset of CD4^+^ and CD8^+^ T-cell blasts from healthy donors expressed high levels of CD27 (**Fig. 6B**). In contrast, CD27 expression on CD4^+^ and CD8^+^ T-cell blasts from P2 and P3 was completely abolished, as confirmed using two distinct anti-CD27 mAbs (**Fig. 6B**). Soluble CD70 binding was also assessed. While subsets of CD4^+^ and CD8^+^ T-cell blasts from a healthy donor exhibited sCD70 binding in a dose-dependent manner, no sCD70 binding was detected in CD4^+^ or CD8^+^ T-cell blasts from P2 and P3. Thus, these results demonstrate that homozygous S70P CD27 abolishes both CD27 surface expression and CD70 binding, confirming its LOF nature in T cells.

**Figure 6.**
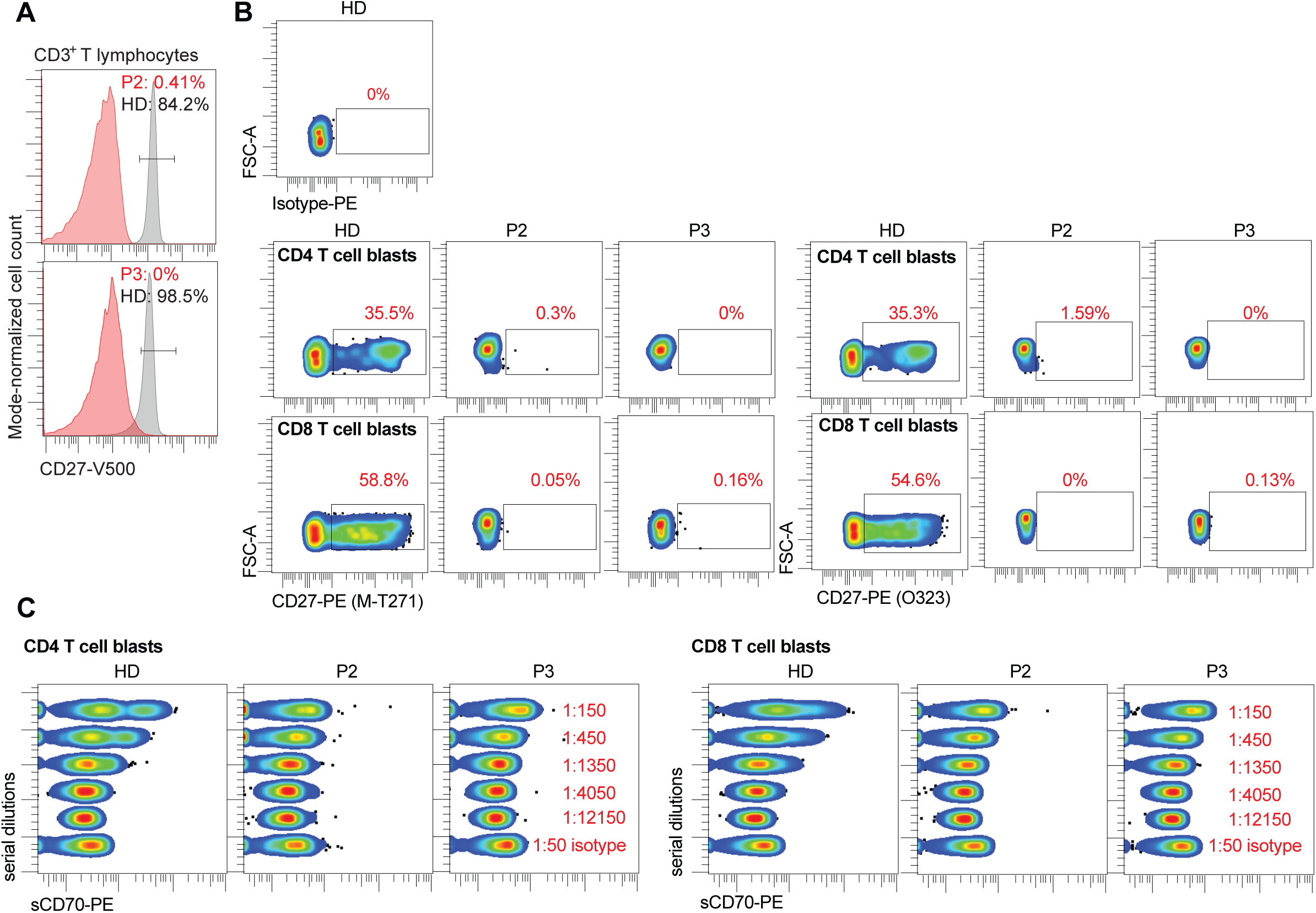
Endogenous S70P CD27 expression is abolished on T cells. **(A)** Surface CD27 expression on CD3^+^ T cells gated from peripheral blood mononuclear cells (PBMCs). **(B)** CD27 expression on CD4^+^ and CD8^+^ T-cell blasts derived from HDs and from patients P2 and P3. **(C)** Soluble CD70 binding on CD4^+^ and CD8^+^ T-cell blasts from HDs and patients P2 and P3, measured across graded concentrations of soluble CD70 or isotype control.

## DISCUSSION

Here, we expand the genotypic spectrum of inherited CD27 deficiency by identifying a novel disease-causing missense variant in the homozygous state in three patients from two unrelated families. This variant was previously reported for P3 in a cohort study, but its pathogenicity was not confirmed (Forbes et al., 2022). This finding underscores the key biochemical role of the Ser70 residue in CD27. Both CD27 and its ligand CD70 belong to the TNF/TNF receptor superfamily, with each forming a homotrimer that engages in a 3:3 stoichiometry. Structural studies have established Ser70 as essential for CD27 function (Liu et al., 2021). First, Ser70 forms two hydrogen bonds with Ile66 of CD27, stabilizing its structure. Second, Ser70 engages Ala80 of CD70 through a hydrogen bond and a van der Waals interaction with Leu81 of CD70 at the CD70–CD27 interface. Third, biochemical assays of soluble CD27 ectodomain carrying the S70A substitution demonstrated slow association kinetics with sCD70 (Liu et al., 2021). Consistent with this biochemical and crystallographic evidence, the S70P variant disrupts both surface expression of CD27 and its ability to bind CD70. These findings also highlight a likely nonredundant role of Ala80 in CD70. Interestingly, four variants predicted to disrupt Ala80 of CD70 are present in the general population (gnomAD v2.1, v3.1, and v4.1), including c.233dup (p.Ala80Serfs*22), c.233del (p.Gly78Alafs*33), c.236del (p.Pro79Glnfs*32), and c.239C>T (p.Ala80Val) (Atkinson et al., 2023; Chen et al., 2022; Karczewski et al., 2020), with a combined global MAF of 1.9 × 10^−5^. Extrapolating from structural data, genetic variants affecting Ser72, Asp74, Arg78, His80, Cys81, Ser83, His86, Asn88, Arg113, Lys115, Thr118, or Glu119 may also perturb CD27 function (Liu et al., 2021). Indeed, substitution of Arg78 with tryptophan (R78W) is an established pathogenic *CD27* variant.

From a population genetics perspective, S70P represents an ancestry-biased pathogenic variant for EBV susceptibility. Consistent with our observation that all three patients are of Hispanic/Latino origin, S70P is enriched exclusively in the Admixed American (AMR) cohort of gnomAD v4.1. Only 11 S70P alleles are reported globally with a global MAF of 6.8 × 10^−6^, 9 of which occur in AMR, where the ethnicity-specific MAF is 1.5 × 10^−4^. This suggests that approximately 1 in 3,300 individuals in AMR carries the S70P variant (Atkinson et al., 2023). Although Latino populations derive from diverse and complex mixtures of indigenous American, European, and Sub-Saharan African ancestries, demographic contractions during the colonial era produced founder effects (Browning et al., 2018; Mooney et al., 2018; Ongaro et al., 2019). It is therefore reasonable to hypothesize that a founder effect might contribute to the elevated S70P frequency in AMR. Notably, CAEBV and extranodal NK/T cell lymphoma, which are both EBV-driven, are most commonly reported among Hispanics, in addition to East Asians (Cohen et al., 2011; Haverkos et al., 2016; Kawada et al., 2023). This observation highlights the importance of future haplotyping and ancestral analyses of the S70P mutation, which may pinpoint a subset of founder populations with particularly high carrier frequencies in AMR, similar to the well-described *FOXN1* LOF founder variant in South Italians (*FOXN1* c.792C>T) and the *RAB27A* LOF founder variant (RAB27A c.244C>T) in Qataris (Adriani et al., 2004; Al-Sulaiman et al., 2020).

The three patients, all homozygous for the same S70P missense variant, illustrate the spectrum of clinical outcomes that can arise from an identical underlying genotype. While all experienced severe forms of EBV disease, underscoring the nonredundant role of the CD27–CD70 pathway in anti-EBV immunity, their courses demonstrate phenotypic heterogeneity and variable expressivity. Initial presentations ranged from overwhelming lymphadenitis and immune dysregulation to HL. Disease sequelae spanned from asymptomatic but persistent EBV viremia to CAEBV requiring HSCT. Beyond EBV-driven disease and its complications, other infections were common, despite being confounded by iatrogenic immunosuppression and HSCT complications in P1 and P2 as well as prematurity in P2. P3, in the absence of profound immunosuppression or HSCT, also experienced two episodes of pneumonia. This observation aligns with a prior series of 49 patients with inherited CD27 and CD70 deficiencies, in which more than half of the patients experienced recurrent infections (Ghosh et al., 2020). Collectively, inherited CD27 deficiency confers a selective susceptibility to EBV, while also predisposing to broader immunodeficiency. All three patients are alive, including P1 and P2 who survived HSCT. Remarkably, only ∼20% of donor chimerism was sufficient to clear EBV infection in P1, highlighting that only a small fraction of CD27-expressing EBV-specific T cells can restore effective anti-EBV immunity.

## METHOD

### Human subjects

The patients and their relatives studied here were living in and followed up at Texas Children’s Hospital, Houston, Texas. The study was approved by and performed in accordance with the requirements of the institutional ethics committee. Informed consent was obtained for the patients, their parents, and healthy donors enrolled in the study. Written informed consent was obtained from all participants. Experiments using samples from human subjects were conducted in accordance with local regulations and with the approval of institutional review board of Baylor College of Medicine.

### Whole-exome sequencing

Trio exome sequencing of P2 was completed clinically (Baylor Genetics, Houston, TX). The method used for exome sequencing of P3 has been described previously (Forbes et al., 2022). In brief, exome capture was performed with HGSC-designed Core (52 Mb) or VCRome 2.1 (42 Mb) reagents (NimbleGen), followed by paired-end sequencing on an Illumina HiSeq 2000 (2×101-bp reads). Reads were aligned to the human reference genome using BWA, PCR duplicates were marked with Picard tools v3.1.1, and variants were called using the GATK best-practice pipeline (base quality-score recalibration and HaplotypeCaller v4.1.4.1). Variants were annotated with SnpEff v4.5.

### Sanger sequencing

Genomic DNA was extracted from pre-HSCT whole-blood samples from the patients and a healthy donor. PCR amplication of the genomic region encompassing S70P variant was performed using forward primer 5’–GAA CAT TCC TCG TGA AGG ACT G–3’ and reverse primer 5’–GCT GTG AGC CTT GAA GAA GG–3’. PCR products were purified and sequenced with capillary electrophoresis (Genewiz, Azenta). Resulting chromatograms were analyzed with 4Peaks software (Nucleobytes).

### Amplicon sequencing

Genomic DNA was extracted from saliva from post-HSCT P1 (DNA Genotek). PCR amplication of the genomic region encompassing S70P variant was performed using forward primer 5’–GAA CAT TCC TCG TGA AGG ACT G–3’ and reverse primer 5’–GCT GTG AGC CTT GAA GAA GG–3’. Amplicon sequencing was performed on a Oxford Nanopore Sequencing (ONT) platform (Plasmidsaurus). Nanopore FASTQ reads were imported into R (ShortRead/Biostrings). Per-read Phred scores (Phred+33) were computed from ASCII quality strings, and read length plus summary quality metrics were calculated. Reads were filtered to ∼260 bp and median Phred ≥30. Strand was assigned using the anchor GTGACCAGCATAG (anchor = “+”; reverse-complement = “−”). The sequence between CGGGGGTC (5′) and CCTTCTCTC (3′) was extracted and unique extracted sequences were tabulated by count.

### Generation of *CD27* delivery vectors

A *CD27* coding sequencing (cds, NM_001242.4)-containing vector was obtained commercially (Genscript). Of note, this WT variant contains H233R (rs2532502) allele, which has a MAF of 0.9907 in gnomAD (Karczewski et al., 2020; Zhang et al., 2018). A59T, P138L, Y32H, W33*, R107C, and S70P variants were generated by site-directed mutagenesis using overlapping primers as follows: A59T (forward 5′–AAG GAC TGT GAC CAG CAT AGA AAG ACT GCT CAG TGT GAT CCT TGC ATA C–3′; reverse 5′–GTA TGC AAG GAT CAC ACT GAG CAG TCT TTC TAT GCT GGT CAC AGT CCT T–3′), P138L (forward 5′–CGG TCG TCT CAG GCC CTG AGC CTA CAC CCT CAG CCC ACC CAC TTA CCT TA–3′; reverse 5′–TAA GGT AAG TGG GTG GGC TGA GGG TGT AGG CTC AGG GCC TGA GAC GAC CG–3′), Y32H (forward 5′–CAA GAG CTG CCC AGA GAG GCA CCA CTG GGC TCA GGG AAA GCT GTG–3′; reverse 5′–CAC AGC TTT CCC TGA GCC CAG TGG TGC CTC TCT GGG CAG CTC TTG–3′), W33* (forward 5′–CTG CCC AGA GAG GCA CTA CTA GGC TCA GGG AAA GCT GTG CTG C–3′; reverse 5′–GCA GCA CAG CTT TCC CTG AGC CTA GTA GTG CCT CTC TGG GCA G–3′), R107C (forward 5′–ACT GCC AAT GCT GAG TGT GCC TGT TGC AAT GGC TGG CAG TGC AGG GAC A–3′; reverse 5′–TGT CCC TGC ACT GCC AGC CAT TGC AAC AGG CAC ACT CAG CAT TGG CAG T–3′), and S70P (forward 5′–ATC CTT GCA TAC CGG GGG TCC CCT TCT CTC CTG ACC ACC AC–3′; reverse 5′–GTG GTG GTC AGG AGA GAA GGG GAC CCC CGG TAT GCA AGG AT–3′). WT and mutant *CD27* alleles were subcloned into AttB-*ACE2* delivery vector, which contains AttB recombinate site and Bxb1 recombinase, to replace *ACE2* using cold fusion or Gibson assembly (Shukla et al., 2024) (Takara Bio and New England Biolabs).

### Overexpression of *CD27* alleles

Overexpression of CD27 alleles, including WT, A59T, P138L, Y32H, W33*, R107C, and S70P alleles, was performed following a previously described (Kamath and Matreyek, 2024). In brief, AttB-*CD27* vectors, either WT or variants, were transfected into landing pad HEK293T cells, followed by AP1903 and puromycin selection. BFP^−^iRFP670^+^ cells represented successfully recombined clones carrying the *CD27* allele of interest. Landing pad cells were cultured in DMEM supplemented with 10% FBS and doxycycline throughout all experiments.

### Flow cytometry of landing pad cells

Detection of CD27 expression on landing pad cells was performed by staining recombined landing pad cells with anti-CD27-PE mAbs, either M-T271 (BioLegend) or O323 (BioLegend), for 30 minutes on ice, followed by data acquisition on an LSRFortessa (Becton Dickinson). Detection of sCD70 binding was performed similarly. In brief, sCD70-PE was obtained commercially (SinoBiological) and was used to stain landing pad cells at indicated dilutions for 30 minutes on ice, followed by data acquisition on LSRFortessa (Becton Dickinson). CD27 expression and sCD70 binding were analyzed within live, single, BFP^−^iRFP670^+^ recombined cells. Isotype-PE control was used for experiments.

### Generation of T-cell blasts

100,000 to 200,000 PBMCs from a healthy donor, pre-HSCT P2 and P3 were plated into 96 well U bot plate in 200 µL in RPMI supplemented with 10% FBS. Cells were stimulated with ImmunoCult human CD3/CD27/CD2 T cell activator (StemCell Technologies) in the presence of IL-2 1000 U/mL (Proleukin, Iovance Biotherapeutics). Cells were expanded for 2 weeks and harvested for experiments.

### Flow cytometry of T-cell blasts

T cell blasts were harvested after 2 weeks of expansion and plated in 96 well V bottom plates. Cells were stained with anti-CD27 anti-CD27-PE mAbs, either M-T271 (BioLegend) or O323 (BioLegend), together with anti-CD4-Pacifi Blue (BioLegend) and anti-CD8a-BV785 (BioLegend) for 30 minutes on ice. T cell blasts were also stained with serial dilutions of sCD70-PE ranging from 1:150 to 1:12,150 (SinoBiological), together with anti-CD4-Pacifi Blue (BioLegend) and anti-CD8a-BV785 (BioLegend) for 30 minutes on ice. Samples were acquired on an LSRFortessa flow cytometer (Becton Dickinson). CD27 expression and sCD70 binding were analyzed within live, single, CD4^+^ and CD8^+^ T cells. Isotype-PE control was used for experiments.

### T, B, NK cell enumeration and immunologlobulin measurements

Enumeration of T, B, and NK cells pre-HSCT was performed by the Clinical Immunodiagnostic Laboratory at Texas Children’s Hospital as a CLIA-certified clinical test. Serum immunoglobulin (IgG, IgA, IgM, and IgE) levels were measured by clinical labs according to standard clinical protocols. Age-adjusted reference ranges were used for interpretation of T, B, NK cell and Ig levels. Serum tetanus, diphtheria, and *S.pneumoniae* antibody levels were measured by clinical labs according to standard clinical protocols.

### Lymphocyte proliferation assay

The lymphocyte proliferation assay (LPA), a clinically certified diagnostic test, was performed by the Clinical Immunodiagnostic Laboratory at Texas Children’s Hospital. Pre-HSCT PBMCs were isolated and seeded at 2 × 10⁵ cells per well in 96-well plates, with or without stimulation. Cells were incubated at 37°C in 5% CO₂ for 3 days in the presence of non-specific mitogens—phytohemagglutinin (PHA), concanavalin A (ConA), and pokeweed mitogen (PWM)—and for 5 days with recall antigens—Candida, tetanus toxoid, and diphtheria toxin. Following incubation, ³H-thymidine (tritiated thymidine) was added to each well and the plates were incubated overnight to allow incorporation into newly synthesized DNA of proliferating cells. The amount of incorporated radioactivity was quantified using a liquid scintillation counter, expressed as counts per minute (CPM) per well. The degree of proliferation was determined by comparing the CPM of stimulated wells to that of unstimulated controls. The stimulation index (SI) was calculated as the ratio of the average CPM of stimulated cells to the average CPM of unstimulated cells.

### NK spontaneous cytotoxicity assay (NKSCC)

NKSCC, a clinically certified diagnostic test, was performed by the Clinical Immunodiagnostic Laboratory at Texas Children’s Hospital. NK cell cytotoxic activity was assessed using pre-HSCT PBMCs isolated from sodium heparin–treated whole blood, both in the presence and absence of IL-2. K562 target cells were labeled with ⁵¹Cr (chromium-51) in complete RPMI medium and plated in 96-well round-bottom plates in triplicate at a concentration of 1 × 10⁴ cells/mL. Effector (E) NK cells derived from PBMCs were added to target (T) cells at E:T ratios of 100:1, 50:1, 25:1, 12.5:1, 6.25:1, and 3.13:1, followed by incubation at 37°C in 5% CO₂ for 4 hours.

Supernatants were harvested, and ⁵¹Cr release was quantified using a gamma counter. Target cells treated with 1% NP-40 represented maximum (total) lysis, whereas target cells incubated without effector cells represented spontaneous release. All measurements were performed in triplicate. The percentage of specific lysis was calculated using the formula: % specific lysis= [experimental cpm - spontaneous cpm) / (maximum cpm – spontaneous cpm)] × 100.

### NK screen by flow cytometry

Detection and phenotyping of NK cells and their subsets were performed using pre-HSCT whole blood samples. A panel of fluorescently labeled monoclonal antibodies was used: CD57–FITC, CD16 (clone 3G8)–PerCP-Cy5.5, CD56–PE-Cy7, CD3–APC-H7, CD45–V450, CD27–V500, NKG2D–PE, and CD16 (clone B73.1)–APC. Following antibody staining, red blood cells were lysed and white blood cells were fixed with paraformaldehyde using the TQ-Prep System (Beckman Coulter). The lysed and fixed cells were washed with BD Pharmingen stain buffer containing BSA, resuspended in the same buffer, and analyzed on a FACS Canto II flow cytometer (Becton Dickinson).

## ACKNOWLEDGMENTS

We thank the patients and their families for their participation in this study; the members of the laboratory for helpful discussions; and Daniela Carrasco Di Lallo, Julia Schauer, Brooke Smith, Shelby Lopez, Adriana Kindelan, and Samantha Johnson for their administrative assistance. We thank physicians and providers, including but not limited to Robert Krance, MD; Manuel Silva Carmona, MD; Stacey Shubert APRN; Nitya Gulati, MD; Nicholas Rider, DO; Kenneth McClain, MD; Ashley Reiland, APRN; Caridad Martinez, MD; Grace Kim, MD; Hari Tunuguntla, MD; and Celine Hanson, MD. We acknowledge the William T. Shearer Center for Human Immunobiology at Texas Children’s Hospital for their equipment facilities, support, and technical assistance in this research.

## SUPPLEMENTARY FIGURE LEGENDS

**Figure 1 Supplement 1.**
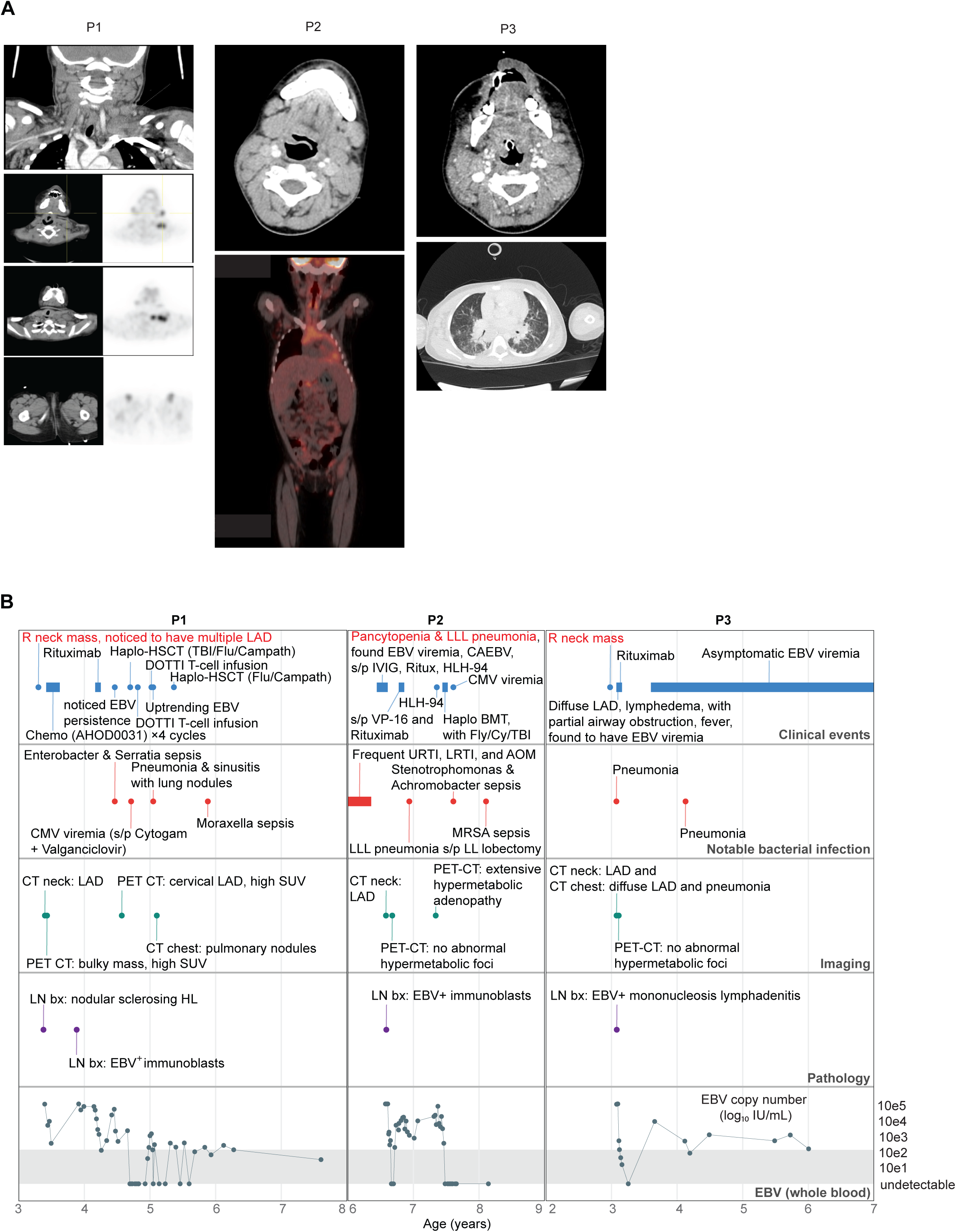
Clinical history of three patients with a homozygous S70P mutation. **(A)** Radiographic and ^16^F-FDG positron emission tomography-computed tomography (PET-CT) images representing metabolically active lymphadenopathy. **(B)** Longitudinal timeline of key clinical events for patients P1, P2, and P3.

**Figure 1 Supplement 2.**
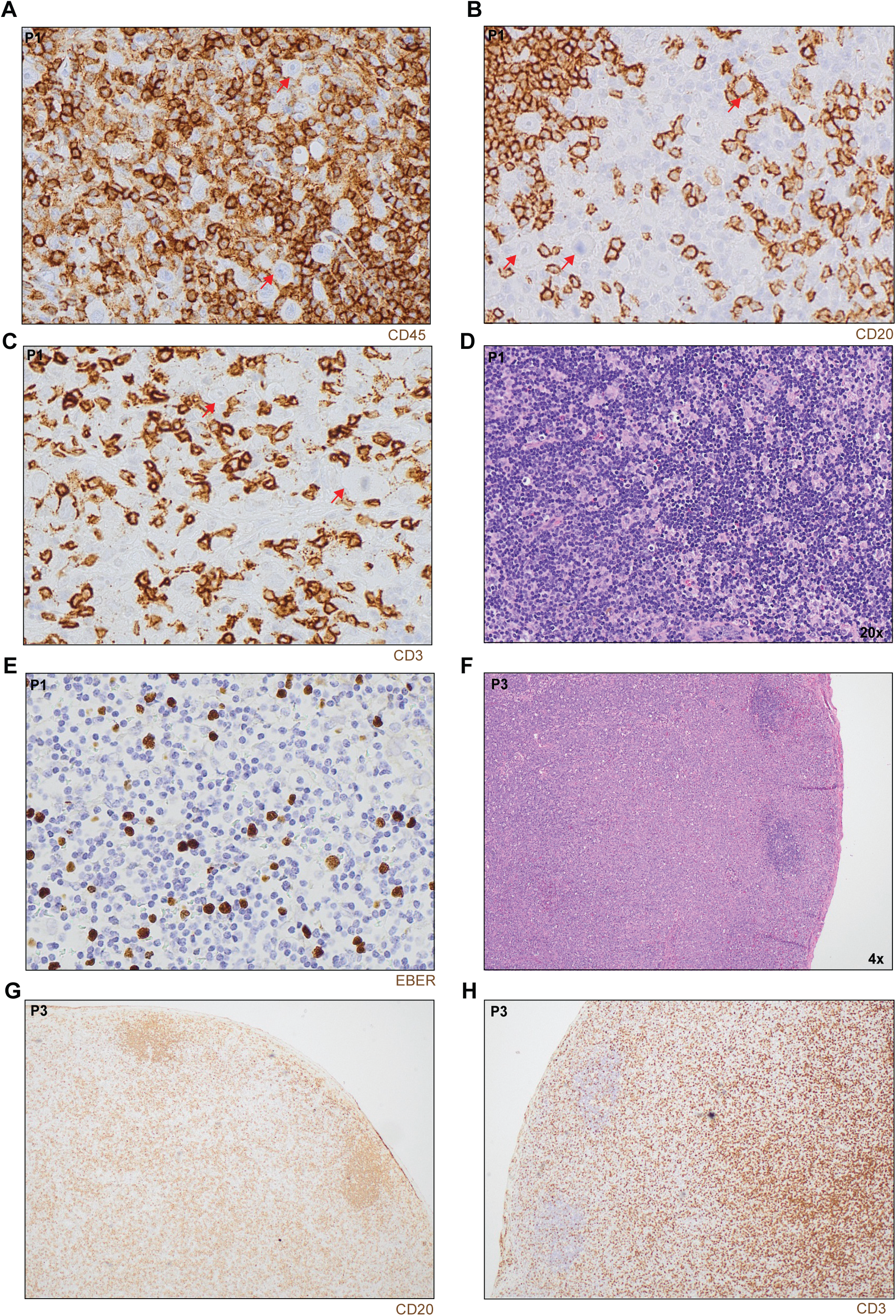
Histological features of lymph node biopsies from P1 and P3. **(A – C)** Immunohistochemistry for CD45 (A), CD20 (B), and CD3 (C) in a lymph node biopsy from P1. **(D)** H&E image (20x) of a lymph node biopsy from P1 after induction chemotherapy. **(E)** EBER in situ hybridization on a lymph node biopsy from P1 after induction chemotherapy. **(F)** Low-power (4x) H&E image of a lymph node biopsy from P3. (**G** and **H**) Immunohistochemistry for CD20 (G) and CD3 (H) in a lymph node biopsy from P3.

